# Family-based Selection: An Efficient Method for Increasing Phenotypic Variability

**DOI:** 10.1101/2025.02.28.640850

**Authors:** Shraddha Lall, Chris Milton, Benjamin de Bivort

**Affiliations:** Department of Organismic and Evolutionary Biology, Harvard University; Center for Brain Science, Harvard University

## Abstract

Persistent idiosyncrasies in behavioral phenotypes have been documented across animal taxa. These individual differences among organisms from the same genotype and reared in identical environments can result in phenotypic variability in the absence of genetic variation. While there is strong evidence to suggest that variability of traits can be heritable and determined by the genotype of an organism, little is known about how selection can specifically shape this heritable variance. Here, we describe a Python-based model of directional artificial selection for increasing the variability of a polygenic trait of interest. Specifically, our model focuses on variability in left-vs-right turn bias in *D. melanogaster*. While the mean value of turn bias for a genotype is non-heritable and constant across genotypes, the variability of turn bias is a heritable and polygenic trait, varying dramatically among different genetic lines. Using our model, we compare different selection regimes and predict selection dynamics at population and genetic levels. We find that introducing population structure via a family-based selection regime can significantly affect selection response. When selection for increased variability is implemented on the basis of independently measured traits of individuals, the response is slower, but leads to a population with a greater genetic diversity. In contrast, when selection is implemented by measuring traits of families with half or full siblings, the response is faster, albeit with a reduced final genetic diversity in the population. Our model provides a useful starting point to study the effect of different selection regimes on any polygenic trait of interest. We can use this model to predict responses of laboratory-based selection experiments and implement feasible experiments for selection of complex polygenic traits in the lab.

## Introduction

Individuals exhibit idiosyncratic behavioral phenotypes, even in the absence of genetic variation [11, 44]. These behavioral differences persist over long periods of time and have been found in behaviors such as activity, aggression [4, 7, 28] and exploration [14, 32, 52], in species such as fruit flies [26], Amazon mollies [3], and mice [18]. At the mechanistic level, such differences may be an unavoidable consequence of the stochastic nature of molecular and cellular function, particularly during development [47, 54]. Clonal single-celled organisms exhibit a significant amount of phenotypic variability, attributed to stochasticity in gene expression [17]. In fruit flies, 23% of genes were differentially expressed among genetically identical individuals reared under essentially identical laboratory conditions [29]. Stochasticity in gene expression may affect behavior through its impact on neuronal physiology, and specific neurons have been implicated in determining specific idiosyncratic behaviors. For example, the degree of left-right asymmetry in the wiring of fruit fly dorsal cluster neurons (DCN) has been shown to predict individual differences in visual object orientation in freely walking flies [30]. Individual differences in calcium responses [24] of specific glomeruli in the antennal lobe of fruit flies predict individual odor preferences [9]. In zebrafish, short-term habituation of the acoustic startle response has been found to be modulated by serotonergic neurons of the dorsal raphe nucleus (DRN) at the individual level [36].

At first glance, the stochastic, noisy nature of molecular processes suggests that intragenotypic behavioral variability may be non-adaptive. But theoretical models [13, 31, 58, 35], and empirical studies from microbes [46, 2] and plants [37, 46, 45, 8] suggest that phenotypic variability can, in some circumstances, be adaptive, allowing organisms to “hedge their bets” against fluctuations in the environment. If variability is adaptive, it may be the object of evolution by natural selection, provided there is heritable variation for the level of variability. Evidence that variability of a trait is under genetic control (and therefore heritable) is mounting, with empirical studies of morphology [12, 16, 51] and behavior [1, 50] supporting this. Quantitative loci whose allelic states impact the variance of a trait, termed ’vQTL’, have been found across microorganisms, plants and animals, including humans [42]. An analysis of the genetic components of variability, both within individual and between individuals, can be found in [22]. These studies suggest that selection can act to increase or decrease variability in trait values. Consequently, bet-hedging strategies can potentially be fine-tuned to the ecological and environmental fluctuations that favor them.

Selection for phenotypic variability has been studied from a quantitative genetics perspective and most extensively applied in plant and animal breeding [57]. Theoretical models predict that when a trait is under directional selection, there are correlated effects on the trait variance [23]. Truncation selection, for example, retains individuals from with phenotypes above or below a certain threshold, i.e., individuals from either of the tail ends of a phenotypic distribution. If sites determining the trait mean also affect its variability, such truncation selection could indirectly select for individuals with greater variability, since they are likely to be over-represented in the tails of the distribution. In animal and plant breeding, the aim is often to increase the mean value of a trait, such as milk production in cows, while reducing variance, to increase uniformity across the population. In such cases, selection can be applied on a combined index that incorporates trait mean and variance, with the aim to increase the former and reduce the latter [10, 34]. Models predict that when the heritability of variance is much lower than mean, selection on such an index will initially result in the mean responding, after which selection pressure will shift to the variance [34]. When disruptive selection is applied on an index of wing shape in fruit flies, there is a drastic increase in phenotypic variation in these traits. Stabilizing and fluctuating selection, however, cause a decrease in phenotypic variation [39]. Selection modeling and experiments have also been applied to within-individual variation, with the aim of reducing variation among repeated measurements made from an individual [25]. These prior studies have not focused on increasing among-individual variability without changing the mean value of the trait.

Demonstrating that variability of a behavioral trait can indeed evolve under artificial selection would be an important step in establishing whether it can evolve in natural circumstances as an adaptive strategy. Locomotor handedness in *Drosophila melanogaster* offers several advantages for such an effort. In fruit flies, locomotor handedness is a behavioral measure that is readily estimated [5] and suitable for experimental evolutionary genetics. Flies have idiosyncratic tendencies to turn left or right in Y-shaped mazes or open arenas [6]. The distribution of average turn biases across flies is unimodal and broad. That is, the most common turn bias is 50-50 left-vs-right, but some individuals have strong left- or right-biases (e.g., in 100 flies, it is common to find individuals with biases as strong as 80-20 or more). Locomotor handedness has a mean of 50-50 across genetic backgrounds [1, 6], indicating that the mean value of turn bias has very low broad sense heritability and is unlikely to respond to selection. In contrast, different wild-derived inbred fruit fly lines exhibit differing degrees of intragenotypic variability in this behavior. This variation has a complex genetic architecture with many sites of small effect correlated with the extent of variability [1]. Consequently, variability of turn bias does have substantial heritability and could potentially respond to selection. Therefore, locomotor handedness has unusual advantages for the study of selection on phenotypic variability, offering little risk that selection on trait variability has confounding effects on the trait mean.

Different approaches can be used over the course of artificial selection to choose which individuals are allowed to mate and contribute to the next generation. In mass selection, also known as individual-based selection, an individual is chosen based on its own score on the trait being selected. Alternatively, selection decisions can be made considering the trait scores of the individual and its relatives (an approach termed family-based or family selection) [55]. The latter approach may enrich the statistical signal in selected individuals, as related individuals may have correlated phenotypes. Additionally, family-based selection can be more likely to maintain allelic combinations that contribute to the selected phenotype non-additively. The families chosen in each generation may be composed of half siblings, often with the same mother but different potential fathers, or of full siblings, generated from one breeding pair. Family-based selection is often employed in plant and animal breeding [49], and is sometimes employed in lab-based selection studies [19, 60].

Laboratory-based selection experiments are long and labor-intensive. Further, factors such as the population size used in the experiment and the number of generations for which selection should be implemented are difficult to predict in advance, and can have significant impacts on the success of the experiment [41]. Theoretical and computational models can be helpful in this regard, allowing one to predict the dynamics of selection at a population and genetic level prior to embarking on a selection experiment. In this paper, we describe an individual-based model of directional artificial selection on a polygenic trait of interest, specifically behavioral variability in locomotor handedness in *Drosophila melanogaster*. Informed by empirical data [1], this model compares the phenotypic and genotypic responses of mass versus family-based selection regimes (both half-sib and full-sib). This study is novel in focusing on a selected trait that is measured using many individuals, the variance of individual behavior. Using this model, we predict how different selection regimes change the underlying genetic architecture of the evolving population, with focus on how epistatic interactions can modulate this change. While our model is motivated by an interest in behavior, it is generalizable and can be readily extended to a diversity of traits with differing genetic architectures. Importantly, it provides specific predictions about responses to selection on a phenotypic and genetic level, helping biologists optimize the parameters of their selection regime before committing to large-scale experiments.

## Methods

Our model assumes a polygenic trait with additive and epistatic components. All individuals are diploid, and reproduction is sexual with a fixed recombination rate. There are no *de novo* mutations. Generations are non-overlapping, with a fixed, finite population size. These assumptions are implemented in the model through variables whose default values are given in Table 1. We chose values appropriate to behavioral experiments in *Drosophila*, but these variables can be readily changed for traits with different architectures or to model populations of various sizes and family compositions.

**Table 1:** Model variables and their default values.

| Variable | Variable Symbol | Default Value |
| --- | --- | --- |
| Population Size | $N$ | 980 |
| Number of Sites | $L$ | 500 |
| Generations | $G$ | 100 |
| Per-base Recombination Rate | $\rho$ | 0.002 |
| Percent Strength of Selection | $s$ | 20% |
| Number of Trials per Individual | $t$ | 500 |
| **For Mass Selection** |  |  |
| Number of Individuals Screened | $N_{sc}$ | $\frac{N}{2}$ |
| Number of Individuals Selected | $N_{sel}$ | $\frac{s}{100} \times N$ |
| **For family-based selection** |  |  |
| Number of Families | $f$ | 35 |
| Number of Individuals per Family | $n$ | $\frac{N}{f}$ |
| Individuals Screened per Family | $n_{sc}$ | $\frac{n}{2}$ |
| Number of Families Selected | $f_{sel}$ | $\frac{s}{100} \times f$ |

The model was implemented entirely in Python 3. Initial code optimization was done on a desktop-PC, and the full model was run on the FAS Research Computing Cluster at Harvard University. Statistical analyses were also performed in Python 3, using built-in numpy [21] and scipy [53] functions. Annotated code can be found at: lab.debivort.org/family-selection-for-variability

We assumed that variability as a trait is determined by *L* sites, each of which has two possible alleles: one that has no effect and one that contributes a direct additive effect and contributes to epistatic effects. Specifically, we assumed that additive effects (*A_i_*) come from a beta prime distribution (Fig. S1), which closely matches the empirical distribution of effect sizes for SNPs associated with variability in locomotor handedness (Ayroles et al., 2015). We used beta prime parameters of *a* = 0.976, *b* = 5.789, loc = 2.79 × 10^−7^ and scale = 0.272. To model epistatic effects, we defined an *L* x *L* symmetrical interaction matrix with each entry *E_i,j_* drawn from an exponential distribution with parameter *λ* and sign randomly assigned as positive or negative with equal probability (i.e., a Laplace distribution with mean 0 and variance 2*λ*^−2^). The value of *λ* was determined based on the desired relative contribution of epistasis to the overall effect size, further detailed below. The values of site effect sizes and the epistatis interaction matrices were drawn once and fixed for all runs and all regimes, thus avoiding any effects introduced by random variation in effect sizes. All individuals in our simulations were diploid, so each site has two loci — individuals could have 0, 1 or 2 alleles at each site with an effect on variability. Thus, the variability trait value for individual *a* was given by

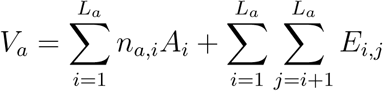

where *L_a_* is the number of sites in individual *a* for which at least one allele has an effect on variability, *i* and *j* index those sites, and *n_a,i_* is, for individual *a*, the number of alleles at site *i* which have an effect on variability (1 or 2 depending whether that site is heterozygous or homozygous). Three levels of epistasis, defined as the ratio of the variance in epistatic effects to the overall variance of genotypic values across the population, were simulated. Specifically, for high epistasis, 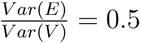, *λ* = 0.0021; for medium epistasis, 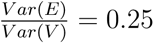, *λ* = 0.0012; for low epistasis, 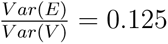, *λ* = 0.00079.

In our model, an individual’s genotype determines their potential to have a phenotype far from the mode [1]. In other words, if an individual comes from a high variability genotype, it will have a phenotype that is drawn from a wider distribution of potential trait values. The particular behavioral trait inspiring this analysis, locomotor handedness, has a mean value very close to 0.5 across all assessed genotypes [5, 6, 1]. Therefore, we used each individual’s *V_a_* value to compute a true locomotor handedness phenotype *P_a_* from a normal distribution with mean 0.5. In order to make the standard deviation for these distributions match empirical data, *V_a_* values were scaled and matched via linear interpolation (using the `interp` function in numpy) to a distribution of empirical standard deviations of locomotor handedness for 167 genotypes (Fig. S2). Accordingly, for each individual, the phenotype was determined as: *P_a_* ∼ *Norm*(0.5, (*d*(*V_a_*))^2^), where *d*(*V_a_*) is the genotypic value after mapping. If the sampled value was less than 0 or greater than 1, we set it to 0 or 1, respectively. Once the true phenotype of an individual was assigned, we could determine its phenotype as would be measured in an actual selection experiment (*π_a_*). Locomotor handedness is measured as the fraction of *t* turns that an animal makes to the right. Thus, *π_a_* ∼ *Binom*(*t, P_a_*)*/t*.

New offspring individuals in our simulations were generated via sexual reproduction. The sex ratio at birth was fixed at 50:50, but the distributions of fitness varied by sex. All females had equal probability of being a parent. However, fruit fly males do not have equal likelihood of successfully mating and fathering offspring. Accordingly, the probability of each male being a parent was drawn from the empirical distribution of male fitness in Fig. 1 of Pischedda and Rice, 2011 [40]. For each new offspring, each of their diploid parents produced two gametes, with recombination. Each site had an independent probability *ρ* of being a site of recombination. For our default values of *L* = 500 sites and recombination rate *ρ* = 0.002, there was on average one recombination event per haploid genome per generation. Of the two gametes generated by this process, one was chosen at random as the haploid contribution to the offspring. Note, in *Drosophila* there is no recombination in males [33], so in this respect our simulation was more typical of other species.

**Figure 1:**
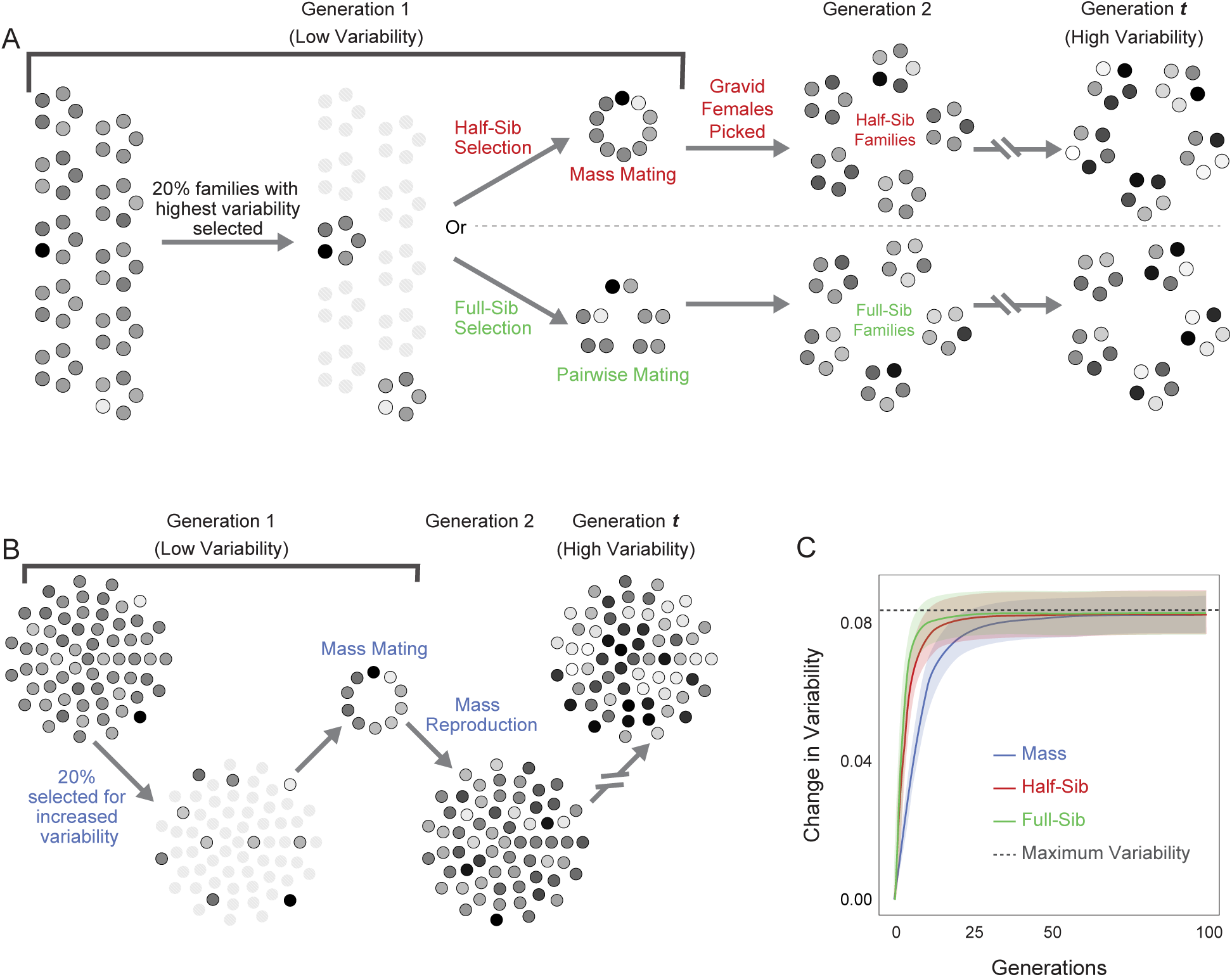
Variability selection regimes. (A) Schematic of family-based selection regime used in the model, showing half-sib and full-sib variants; (B) Mass selection regime; (C) Change in turn bias variability (Δ*σ_g_* = *σ_g_* − *σ*_0_) over 100 generations of mass, half-sib and full-sib selection. Lines are the average Δ*σ_g_* at each generation *g*, averaged across *k* = 10000 replicate runs of model. Shaded regions indicate ±1 standard deviation due to run-to-run variation (*ɛ_r_*)*_g_* estimated as the standard deviation of Δ*σ_g_* across *k* replicates; dotted line indicates the variability value of a population where all individuals have the maximum variability trait value *V_a_* possible with this genetic architecture.

In every generation of both family-based and mass selection simulations, we implemented a strength of selection *s* = 0.2 or 20%. For simulations of family-based selection, the initial population was seeded with *f* families, each of which have *n* individuals, for a total population size of *f* ∗ *n* = *N* . The default values of *f* = 35 and *N* = 980 were chosen as plausible numbers for a potential real implementation of this experiment. Measured phenotypes (*π_a_*) were computed for all of the individuals from each family. The measured variability for each family *m*, in generation 0, was calculated as the standard deviation of individual phenotypes: *σ_m_*(0) = *std*(*π_a_*). *s*% of *f* families with the highest variability were selected to seed the next generation (Fig. 1A). The average population variability was estimated as the average of family variabilities for all *f* families: *σ*_0_ = *µ*(*σ_m_*(0))

For half-sib selection, individuals from all selected families were pooled, and *f* families were seeded for the next generation by randomly picking *f* multiply-mated females from the pool. For full-sib selection, male and female individuals were randomly paired from among the selected families to create *f*monogamous pairs, that were used to generate *f*families for the next generation. The variability for all families in the population (*σ_m_*(*g*)) was estimated at every generation *g*, and selection was carried out as before.

For simulations of mass selection, the initial population was seeded with *N* individuals. Measured phenotypes (*π_a_*) were computed for all of the individuals in the population. The variability of this population was calculated as the standard deviation of the measured phenotypes: *σ*_0_ = *std*(*π_a_*). *s*% of *N* individuals were chosen to seed the next generation, such that the selected individuals formed a subset with the highest possible variability (Fig. 1B). This was implemented by sampling across the distribution of individuals, selecting those that have the highest probability of being from a target high variability distribution. This was the default method we used to implement mass selection, and we refer to this method of mass selection as ‘uniform sampling.’ We additionally implemented a mass selection approach we term ‘extreme sampling’, wherein the 10% of the individuals with the most extreme left-bias and the 10% with the most extreme right-bias were selected every generation. This is a form of two-tailed truncation selection. In both cases, the next generation of *N* individuals was created via reproduction among all selected individuals. The variability of the population *σ_g_* was estimated at each generation, and selection was carried out as before.

We estimated the selection differential (*S*) in different regimes by calculating the difference in variability of the whole population and variability of the subset of individuals that was selected every generation. Selection response (*R*) was calculated as the change in phenotypic variability over 1 generation of selection. We quantified the realized heritability (*h*^2^) at different time points during selection as the slope of the regression between the cumulative changes in *S* and *R*. At generation *g*, 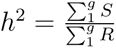

To track the genotypic changes associated with selection, each variability site was evaluated for fixation at the end of selection. All selection regimes with various combinations of parameters were simulated *k* times under different random seeds to assess variation in selection response trajectories.

## Results

### Family-based selection for variability is faster than mass selection

Starting with a simple additive model (no epistasis), our simulations showed robust responses to selection for higher turn bias variability (Fig. 1C). Family-based selection (half-sib or full-sib) yielded a faster response than mass selection. Half-sib family-based selection yielded 75% of its total response to selection by generation 6, whereas mass selection attained 75% of its final response by generation 11. Full-sib selection was slightly faster than half-sib selection, attaining 75% of its final response approximately 2 generations sooner. When we implemented mass selection with extreme sampling, the response was faster than with uniform mass selection, but was still slower than both half and full-sib family-based selection through at least 5 generations (Fig. S3). However, a downside of extreme sampling is the possibility of evolving an undesirable multimodal phenotype distribution if the mean of the trait is also heritable (Fig. S10).

Initial populations had comparable variability (*σ*_0_ = 0.135)across regimes, measured as the standard deviation of phenotypes across all individuals. The change in variability over 100 generations of selection was consistent across all regimes. The final variability attained was, on average, almost as high as the theoretical maximum possible value, calculated as the variability of a population where all the individuals have all the sites fixed at the high variability allele (Fig. 1C).

We estimated the selection differential *S* across regimes as a function of the number of generations of selection. Mass was associated with the highest selection differential at all generations (Fig. S4A). But that regime’s selection response (*R*) was only slightly higher than that of the family-based regimes from generations 6-15, and even there only by a small amount (Fig. S4B). Mass selection moreover had a much lower selection response than family-based regimes for the first 3 or 4 generations. Thus, the realized heritability of family-based selection was higher than that of mass selection consistently across generations.(Fig. S4C).

### Family-based selection for variability fixes more sites than mass selection

Many more variability-determining sites were fixed in 100 generations of family-based selection compared to mass selection, with full-sib selection fixing 94.9% of all sites on average (Fig. 2A). This was an expected consequence of the lower effective population size, and therefore higher inbreeding in family-based selection [38]. Sites with large effects on turn bias variability had higher fixation probabilities in all regimes (Fig. 2B-D), as expected. This relationship was strongest in mass selection (*r*^2^ = 0.81 using a second-degree polynomial fit), weaker in half-sib selection (*r*^2^ = 0.20), and weakest in full-sib selection (*r*^2^ = 0.07), where a large number of sites were likely to fix regardless of their effect size (Fig. 2D).

**Figure 2:**
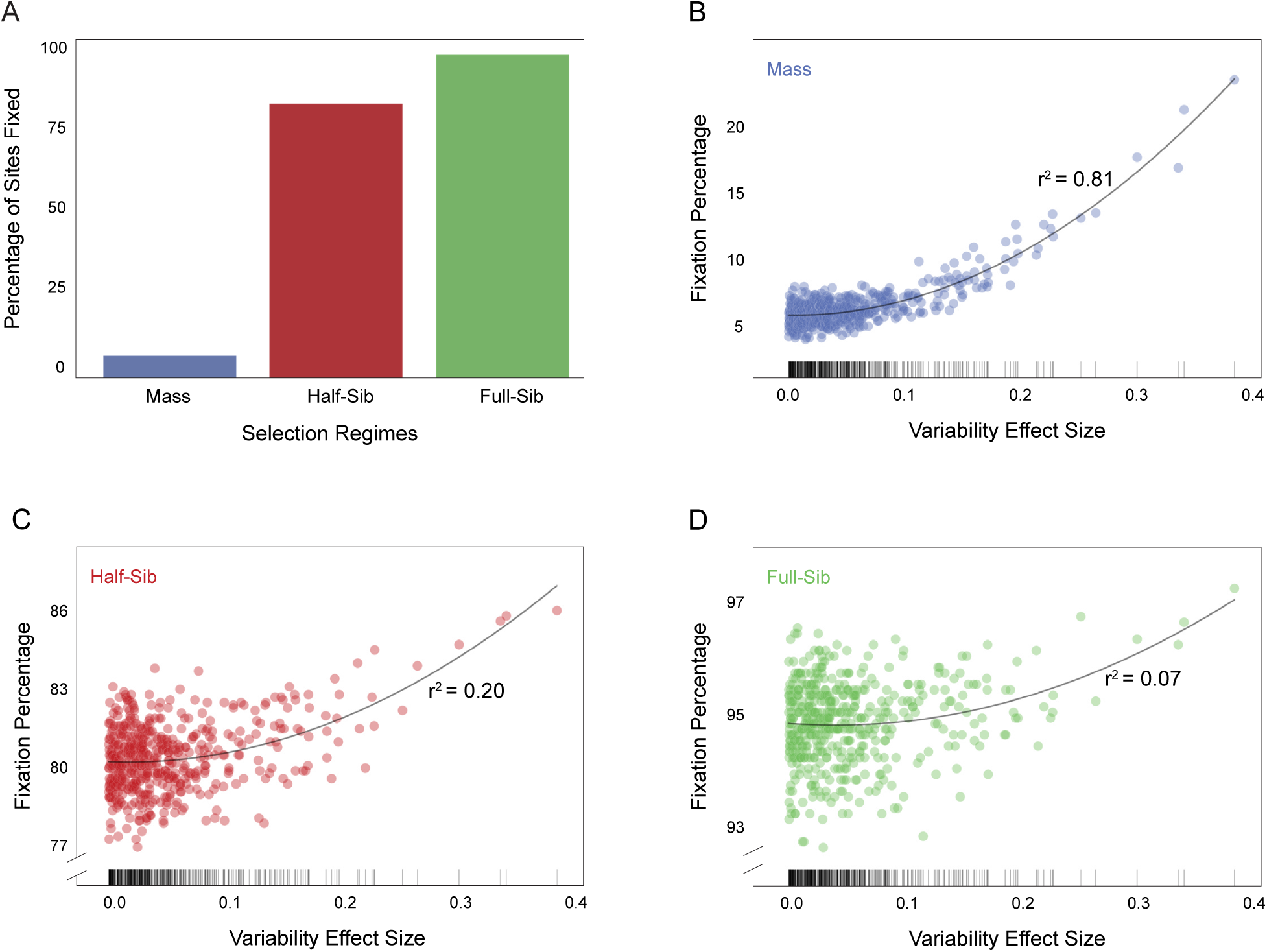
Fixation of variability determining sites after 100 generations of selection. (A) Mean fixation percentage of sites at generation 100 for the three selection regimes, averaged over 1000 runs. (B-D) Site fixation probability versus effect size in mass, half-sib and full-sib regimes, respectively. Lines are second-degree polynomial fits. Tick marks at bottom indicate the values of site effect sizes, which were fixed across all runs.

### Family structure modulates selection response

To determine the effects of family and population size on selection response, we systematically varied these parameters (*f* and *N* ) and measured peak variability change Δ*σ*_100_ = *σ*_100_ − *σ*_0_. Selection response was smaller at both extremes of *f* (Fig. 3A). The smallest response to selection was found with populations with few families of many individuals each (low *f* , high *n*). Slightly higher variability was attained with populations made of a large number of families with few individuals in each family (high *f* , low *n*). The greatest selection responses were found in populations with families of intermediate size. Regardless of family composition, increasing population size boosted the selection response.

**Figure 3:**
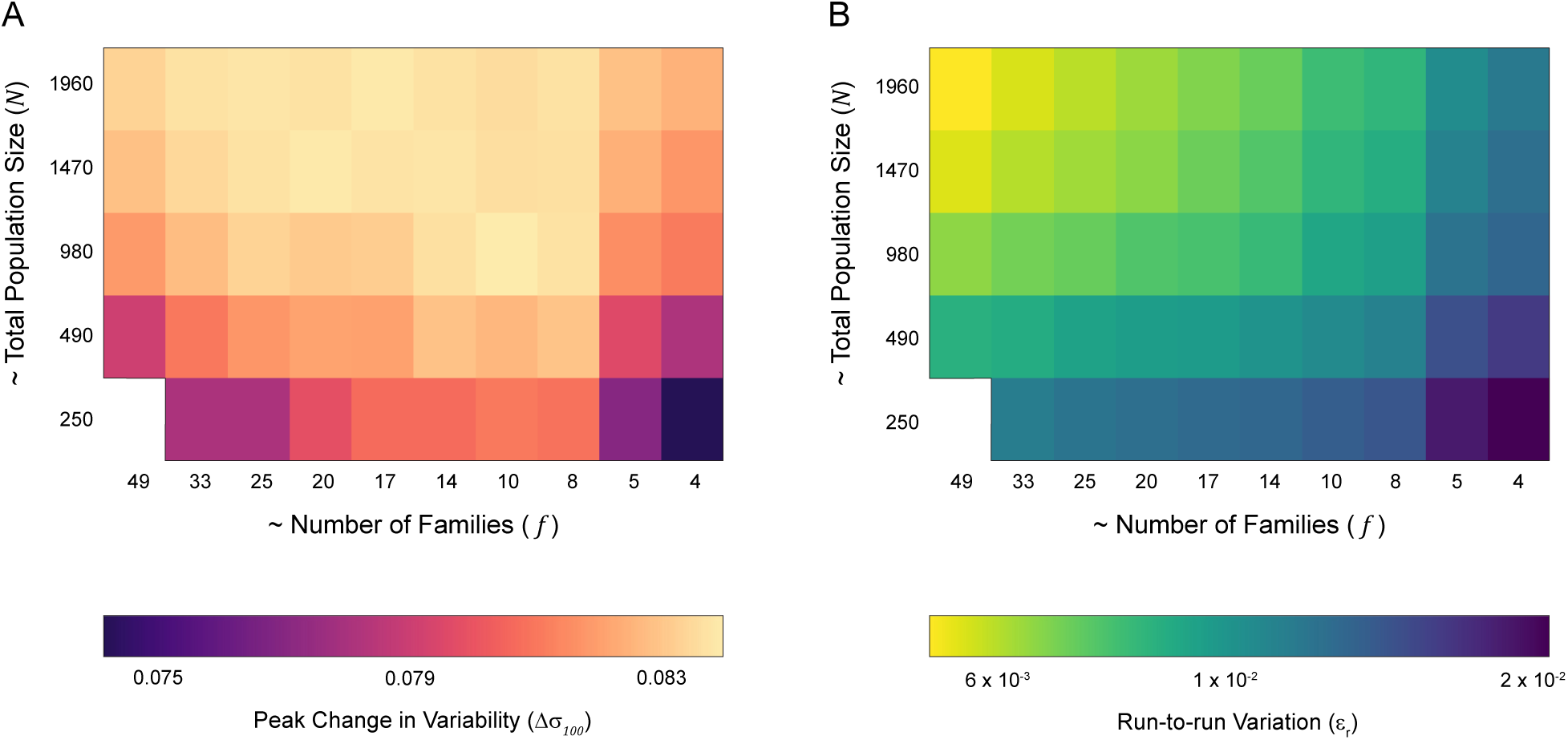
Effects of population structure in family-based selection. Changes in (A) peak variability (Δ*σ*_100_) and (B) run-to-run variation (*ɛ_r_*) in selection response over 100 generations of half-sib selection. Each row has a fixed total population size, with population structure varied by changing the distribution of animals within families. The leftmost column corresponds to a many families with few individuals in each, while the rightmost column corresponds to fewer, larger families.

Run-to-run noise (*ɛ_r_*) can affect the reliability of selection, an important consideration for artificial selection experiments that might actually be implemented in the lab. With fewer, larger, families (low *f* , high *n*), selection responses were more variable (Fig. 3B). Populations with more, small families (high *f* , low *n*) were generally less subject to noisy fluctuations. Unsurprisingly, increasing total population size reduced run-to-run variability. Overall, our simulations indicated that intermediate number of moderate sized families can optimize selection for variability at various population sizes, allowing for reliable, consistently high responses.

### Selection Outcomes are Robust to Change in Model Parameters

Changing model parameters (in Table 1) altered the selection responses quantitatively, but did not alter the overall trends and differences between the selection regimes. For example, changing the number of sites that contribute to phenotypic variability from 63 to 1000 had little discernible effect on the selection response in any of the regimes (Fig. S5). Likewise, changing the distribution of site effects had no qualitative effect on selection outcomes (Fig. S6). This suggest that results from our model apply to traits with a variety of effect size statistics.

When selection was weaker (i.e., more individuals were chosen to found subsequent generations) the selection response was slower, as expected. This effect was most pronounced under mass selection (Fig. S7). Modulating population size had similar effects across all regimes, with larger population sizes showing faster and stronger responses to selection. This effect was more pronounced in family-based selection, where the effective population size is reduced due to inbreeding (Fig. S8).

### Epistasis Attenuates Selection Responses

Our model accommodates pairwise epistatic interactions between sites that additively contribute to variability. These epistatic effects can be positive or negative (Fig. 5B), increasing or decreasing an individual’s latent variability (*V_a_*). When epistasis was added to the model, the response to selection was slower across all selection regimes. However, over 100 generations, a similar value of variability was attained (Fig. 4A). The effect of epistasis was quantitative: increasing the magnitude of epistatic interactions resulted in slower selection responses (Fig. 4B, Supplementary Figure 9).

**Figure 4:**
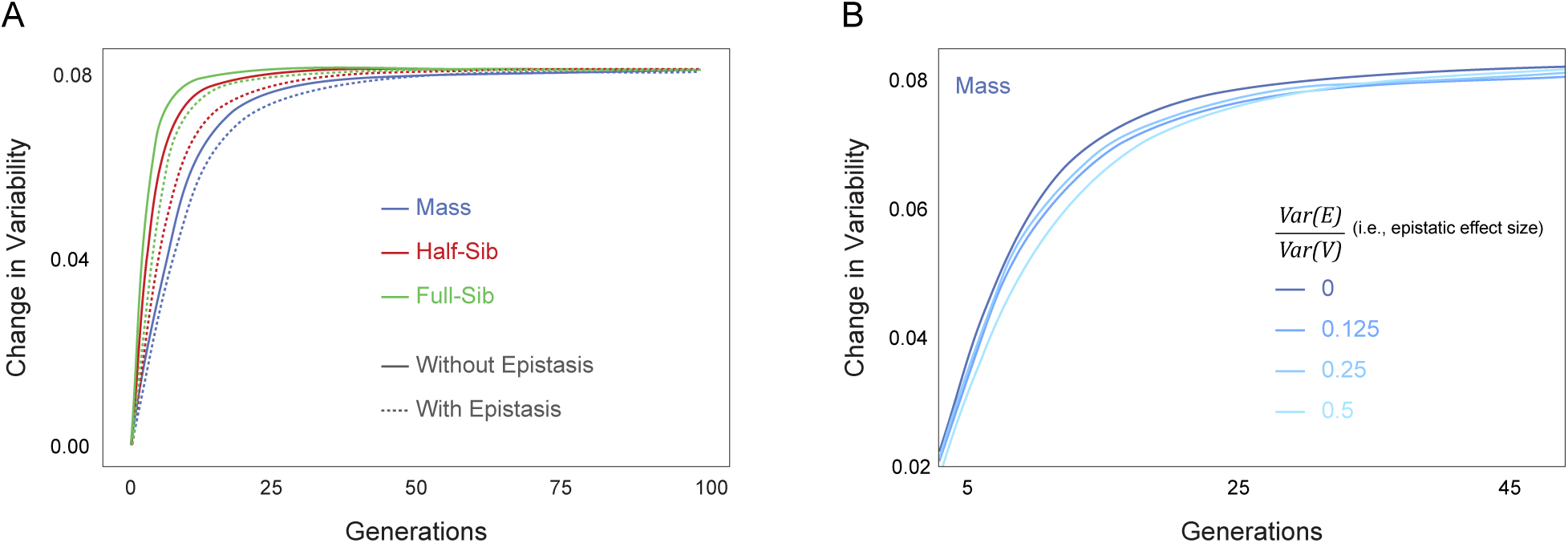
Effect of epistasis on selection response. (A) Average change in variability versus generations of selection, with high level of epistasis 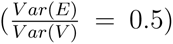 and without epistasis, in different selection regimes; (B) Average change in variability versus generations of mass selection, with varying magnitudes of epistatic interactions.

**Figure 5:**
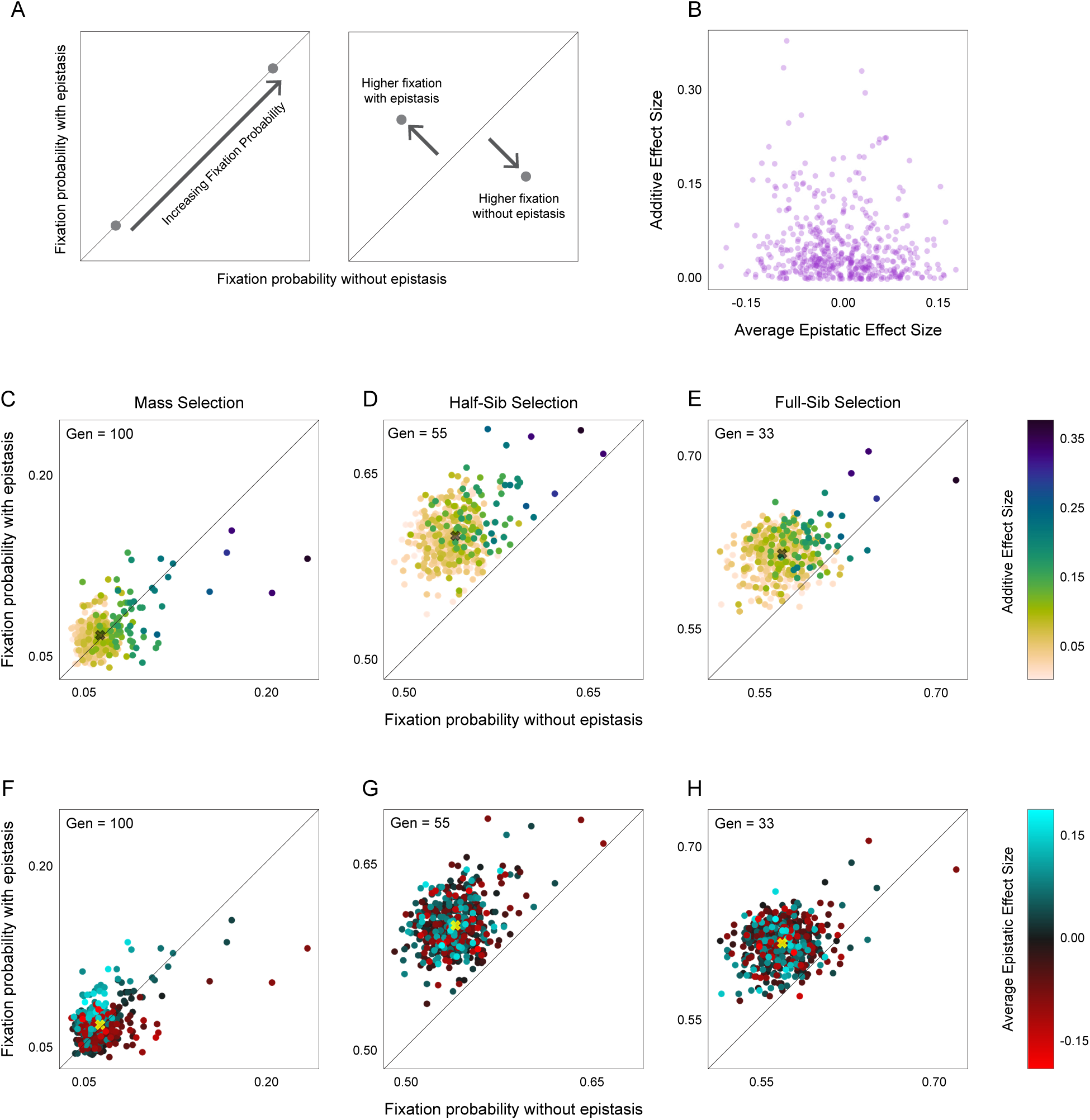
Effect of epistasis on fixation of variability-influencing sites. (A) Schematic for interpreting fixation probability plots in (C to H); (B) Additive effect size versus average epistatic effect size for each site (purple point). (C-E) Site fixation probability with epistasis versus without epistasis, under mass, half-sib and full-sib selection regimes after 100 generations. Points (sites) are colored by their additive effect sizes. Diagonal line indicates the unity line and X marks average fixation probabilities across sites. (F-H) as in C-E, but with points colored by their average epistatic effect.

### Sites Are Differentially Fixed With and Without Epistasis

We examined the effect of epistasis on the probability of site fixation after selection (Fig. 5A). In our model, a site’s additive effect and its average epistatic effect were uncorrelated(Fig. 5B). How these two effects determined a site’s fixation probability depended on the selection regime. In general, fixation probabilities were higher when both additive and epistatic effects were included (Fig. 5C-H). This difference was more pronounced in family-selection regimes, where the population structure could preserve combinations of linked sites with strong epistatic interactions. Unsurprisingly, the fixation probability of a site depended on its additive effect, with larger effect sites being more likely to fix (Fig. 2C-E). This relationship was strongest when epistasis was not included in the model.

Conversely, sites that had very positive average epistatic effects were more likely to fix when the model included epistatis (Fig. 5F-H). This was true even for sites with low additive effects, on account of their large positive epistatic contribution to variability. Similarly, sites that had highly negative epistatic effect sizes fixed with a higher probability with epistasis, potentially due to their strong negative effects on variability driving them to extinction (Fig. 5F-H).

### Family-based Selection is Particularly Efficient for Variance-but not Mean-Based Traits

In our model, family-based selection produced a faster response than mass selection when the trait under selection was variance-based, such as turn bias variability (Fig. 1C, Fig. 6A). We hypothesized that the efficiency of family-based selection resulted from the concordance of the unit of selection (families) and a trait that can only be measured from multiple individuals (variance). To test this hypothesis, we used the same model (genetic architecture, selection regimes, etc.) to instead select for increased mean value of the trait. When the trait under selection was mean-based, and not variance-based, all selection regimes were equally rapid (Fig. 6B). Both mass and half-sib selection yielded close to their maximum selective response within 8 generations (Fig. 6D & F). This was in contrast to what we saw for a variance-based trait, in which mass selection did not yield nearly half of its selective response within 8 generations (Fig. 6C & E), while half-sib selection reached close to maximum variability. Therefore, the increased efficiency of family-based designs appears specific to selection that acts on the variability of a trait, and not its mean.

**Figure 6:**
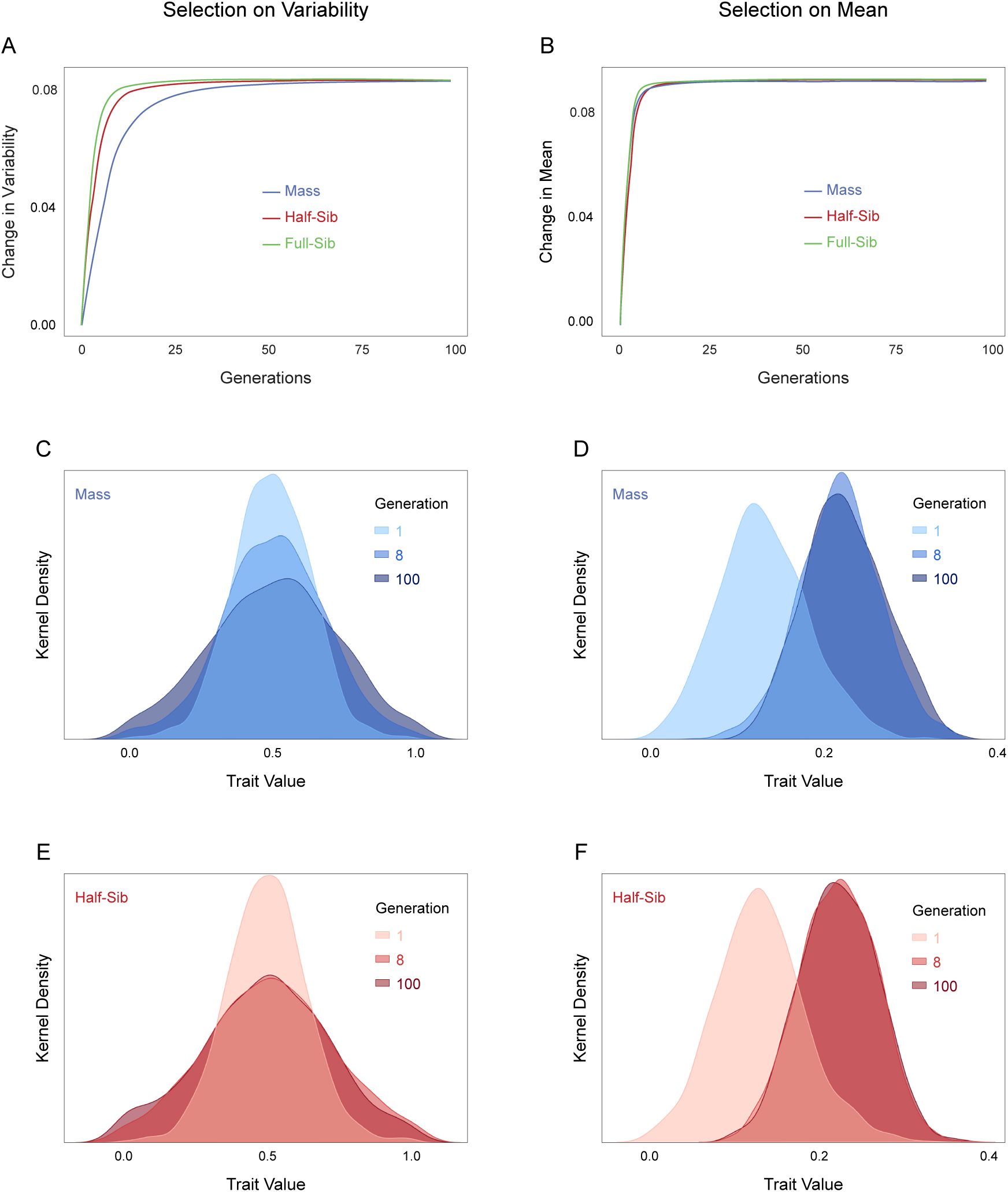
Efficacy of selection regimes for variance-based and mean-based traits. Change in selected phenotype versus generations of selection for increased (A) variability of the trait (same as Fig. 1C) and (B) mean of the trait. (C) Kernel density estimates of the distributions of trait values at generations 0, 8 and 100 of mass selection for increased trait variability. The generation 8 distribution is not as wide as the generation 100 distribution. (D) As in C, but for selection for increased trait mean. Note that the generation 8 distribution has a similar mean to that of generation 100. (E, F) As in C and D, but for half-sib selection. In both panels, the generation 8 and 100 distributions are very similar.

We conducted further analyses of traits that have heritability for both mean and variability. This is a complex topic, with interactions between mean and variability, their respective trait architectures and the selection regime. We explored these relationships only superficially, with a toy trait where mean and variability were determined by independent sites. We used the same family-based and mass regimes to select for increased variability. When variability and mean are both determined by the same number of sites with effect sizes drawn from the same distribution, the overall qualitative relationships remain as before - family selection outperforms extreme and uniform mass selection, especially in the initial generations (Fig. S10A-C). However, selection outcomes can differ when the mean and variability have different underlying effect sizes. When the mean value of the trait has sites of much larger effects than the variability sites (i.e, the mean has higher heritability), selecting on variability is more effective with mass selection (Fig. S10D). As suspected, using extreme (two-tailed truncated) mass selection resulted in the final population having a tri-modal or bi-modal distribution of phenotypes (Fig. S10F).

## Discussion

We examined the efficiency of different artificial selection regimes at evolving variability of a trait. We modeled a trait with a polygenic architecture, including the possibility of epistatic interactions, as is typical of behaviors. Our simulations demonstrated that selection can indeed increase the variability of a polygenic trait (Fig. 1C), without impacting the mean (Fig. 6A,C,E). Family-based selection allowed for an efficient response of increasing variability in the focal trait. The parameters of our trait were based on locomotor handedness in *Drosophila melanogaster*, but should generalize to many traits, and our major findings are robust to large parameter changes (Fig. S5, Fig. S6, Fig. S7, Fig. S8). Mass selection did result in a strong selection response, but this response was slower, a difference with practical consequences for actual selection experiments. For instance, laboratory artificial selection in fruit flies over a year would result in 20 to 25 generations [48]; at this point family-based selection is still likely to have yielded a larger response than mass selection. An added practical advantage of family-based selection is that family-wise variability estimates can be obtained by screening a subset of the individuals in each family [55]. Therefore, a comparable strength of selection can be achieved by screening fewer animals than in mass selection.

Since estimating variance requires the measurement of multiple individuals, it makes sense that family-based selection will be more efficient at selecting for variability. Family-based selection measures the phenotypes of multiple, related individuals that are enriched for common variants, and propagates families as the unit of selection. Consistent with this reasoning, we found that family-based selection was faster than mass selection on our variance-based trait (Fig. 6). Consistently, family-based selection had higher realized heritability (Fig. S4C) despite having lower selection differentials (Fig. S4A). We believe this performance can be attributed to family-based selection estimating the true genetic trait value *V* more faithfully by calculating it over genetically related individuals.

When the same model was used to select for an increase in the mean value of the phenotype, all three regimes performed essentially identically (Fig. 6B,D,E). Therefore, family-based selection might be specifically effective when the selected trait reflects variability among individuals. While turn bias in flies does not have substantial heritability for its mean [1, 5], our model allows the study of selection for variability in traits with both heritable mean and variability. Selection responses can vary depending on how much the mean and variability of the trait vary, their heritabilities, their underlying site effect size distributions (Fig. S4), and almost certainly any pleiotropy between these traits. Our model offers a flexible tool for exploring these relationships, and can help pinpoint the best selection regime for specific traits. Follow-up studies using this model also have the potential to clarify the general relationships between trait mean and variability genetic architectures and efficient selection regimes.

Population size plays a key role in selection, with larger populations being more resistant to drift, resulting in stronger selection response [27, 59]. Accordingly, we found larger population sizes resulted in faster, more reliable responses across all selection regimes (Fig. S8). Family-based selection schemes have smaller effective populations than mass selection [38, 56], and we found that their maximum selection responses were correspondingly sensitive to decreased total population size(Fig. S8). The relative size of families within the population also mattered for selective responses. Given a constant total population size, more, smaller families improved the reliability of selection across independent runs (Fig. 3B). Reliability is important when planning selection experiments in organisms with substantial generation times. Even fruit flies, with 10 day generations, require months or years to complete an artificial selection experiment. Unfortunately, while increasing the number of families improves reliability, it tends to diminish the magnitude of selection responses (Fig. 3A). Therefore, a population that has an intermediate number of moderate sized families represented a good compromise between maximizing selection response and minimizing run-to-run noise. Our model thus is helpful for planning experiments and predicting the population structure most likely to maximize the chances of seeing a selection response within experimental constraints. Laboratory-based evolution experiments in *Drosophila* can show a rapid response. Directional selection on the mean value of thermal tolerance, for example, led to selected flies having a higher thermal tolerance within ten generations [20]. In another study, flies selected for better foraging found food twice as fast as control flies after five generations of selection [43]. In male flies selected for increased or decreased sexual aggression, lineages diverged in the aggressive behavior of forced copulation in 3 to 5 generations [15]. While experimental evolution on variability has not been conducted in flies, these previous findings suggested that our model’s prediction of selection responses within ∼10 generations are plausible. However, there are factors which our model does not account for that might slow the response, such as fitness trade-offs or physiological limitations. Our model considers sites which only affect our trait of interest, but there might be pleiotropic interactions with other traits that could alter the selection response.

Site fixation probabilities varied considerably across regimes (Fig. 2) and with the presence or absence of epistasis (Fig. 5). Even with a large total population size, family-based selection’s reduced effective population size led to substantially lower heterozygosity after selection in our model (Fig. 2A). The effects of inbreeding were most prominent in full-sib selection. The absence of inbreeding in mass selection resulted in far fewer sites being fixed after 100 generations (Fig. 2A). Also, in mass selection the effect sizes of the sites were more strongly correlated with fixation probability. Sites of large effect had higher chances of fixing in all regimes, but sites of low effect were much less likely to fix in mass selection (Fig. 2), consistent with drift being a powerful dynamic in family-based selection. Therefore, while the population structure in family-based selection can accelerate the speed of selection, the loss of genetic diversity could potentially lead to inbreeding depression ([38]).

When epistatis was factored into the model, fixation probability rose (Fig. 5). Since inbreeding and family structure can preserve larger regions of linked sites, they make it more likely that sites with neutral or mild direct effects but more positive or negative epistatic effects will fix (Fig. 5F-H). Conversely, sites of high direct effect can be prevented from fixing if they are epistatically coupled with sites that negatively impact the trait under selection. This was evident in our model, with sites of high additive effect size having a lower likelihood of fixation when epistasis was included, across all selection regimes (Fig. 5C-E). When planning actual selection experiments, the true level of epistasis may be unknown.

Fortunately, our model suggests that the extent of epistasis does not change the relative efficacy of the three regimes (nor did it impact the final variability values attained after many generations of selection; Fig. 4).

Our model is flexible, accommodating populations of varying structures and traits having different underlying genetic architectures. We did not include dominance interactions in our model, but the model could be readily extended to incorporate a dominance matrix, similar to our implementation of the site-wise epistatic matrix. We observed a robust response to selection for variability across a wide range of configurations with our model (Fig. S5 - Fig. S8). Our model highlights that selection can change the variance of traits, and that family-based regimes are an effective way of implementing directional selection for increased variability. Intragenotypic behavioral variability can be a target of selection, in the lab and potentially in nature. Responsiveness to selection is a precondition for bet-hedging to evolve as an adaptive phenotypic strategy. Experimental studies that implement selection on variability in the lab can are a promising path forward to understand the evolutionary role of variability and its genetic basis.

## Data Availability

Annotated code used to generate all the data can be found at: lab.debivort.org/family-selection-for-variability. Empirical data for turn-bias variability in *Drosophila melanogaster* were obtained from [1]. Relevant data can be found at: https://lab.debivort.org/genetic-control-of-phenotypic-variability.

## Acknowledgments

We thank Ian Dworkin for the conceptual understanding of family-based selection, Danylo Lavrentovich for assistance in using the FAS Research Computing Cluster, and Vikram Ravindranath for help with LaTeX typesetting. B.d.B was supported by NIH/NINDS grant no. 1R01NS121874-01.

## Supplementary Figures

**Figure S1:**
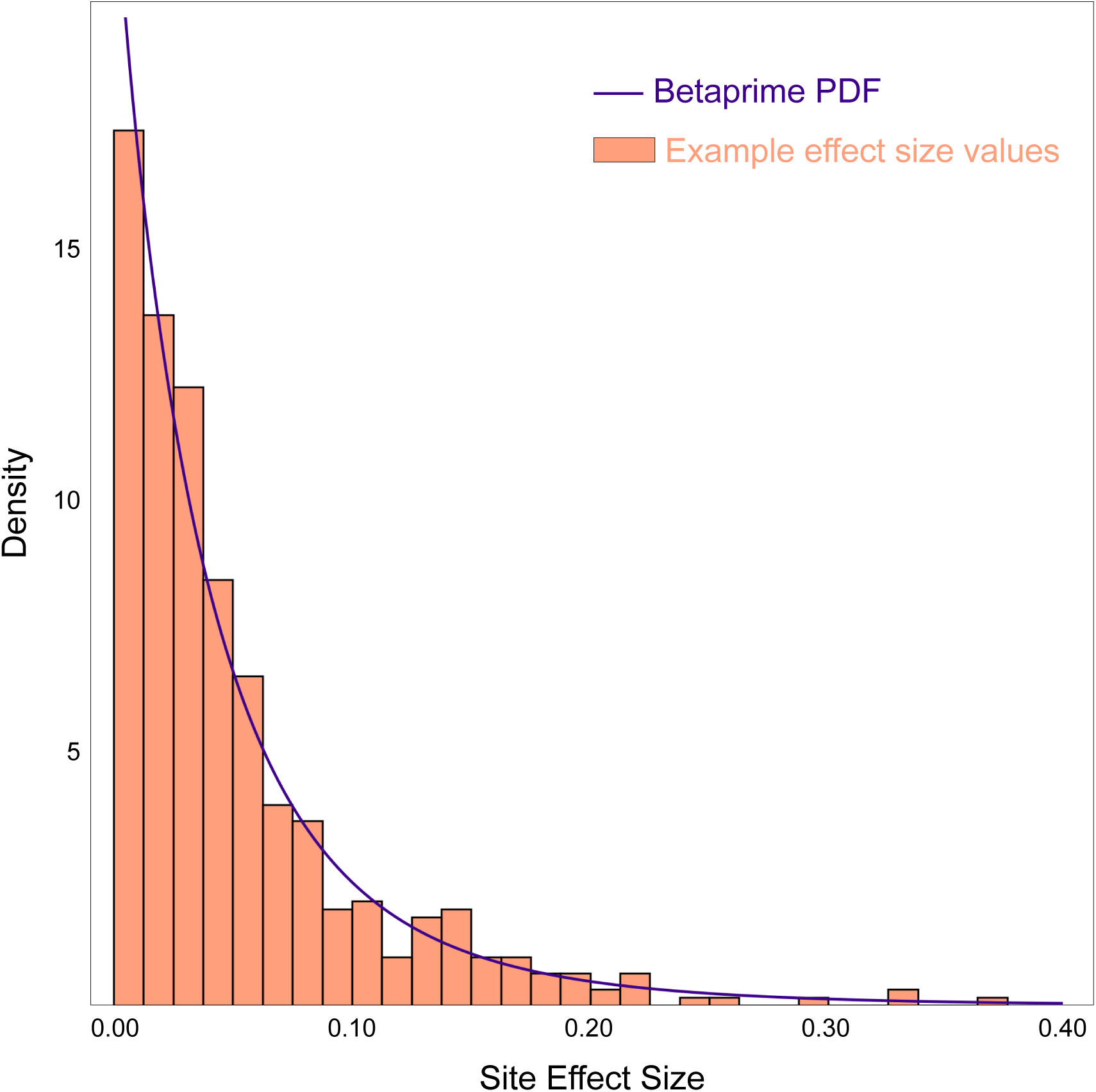
Distribution of Site Effect Sizes. Probability density function of betaprime distribution [1] from which site effect sizes are drawn (black line); example values of 500 sites with effect sizes drawn from the distribution from one particular run of our model.

**Figure S2:**
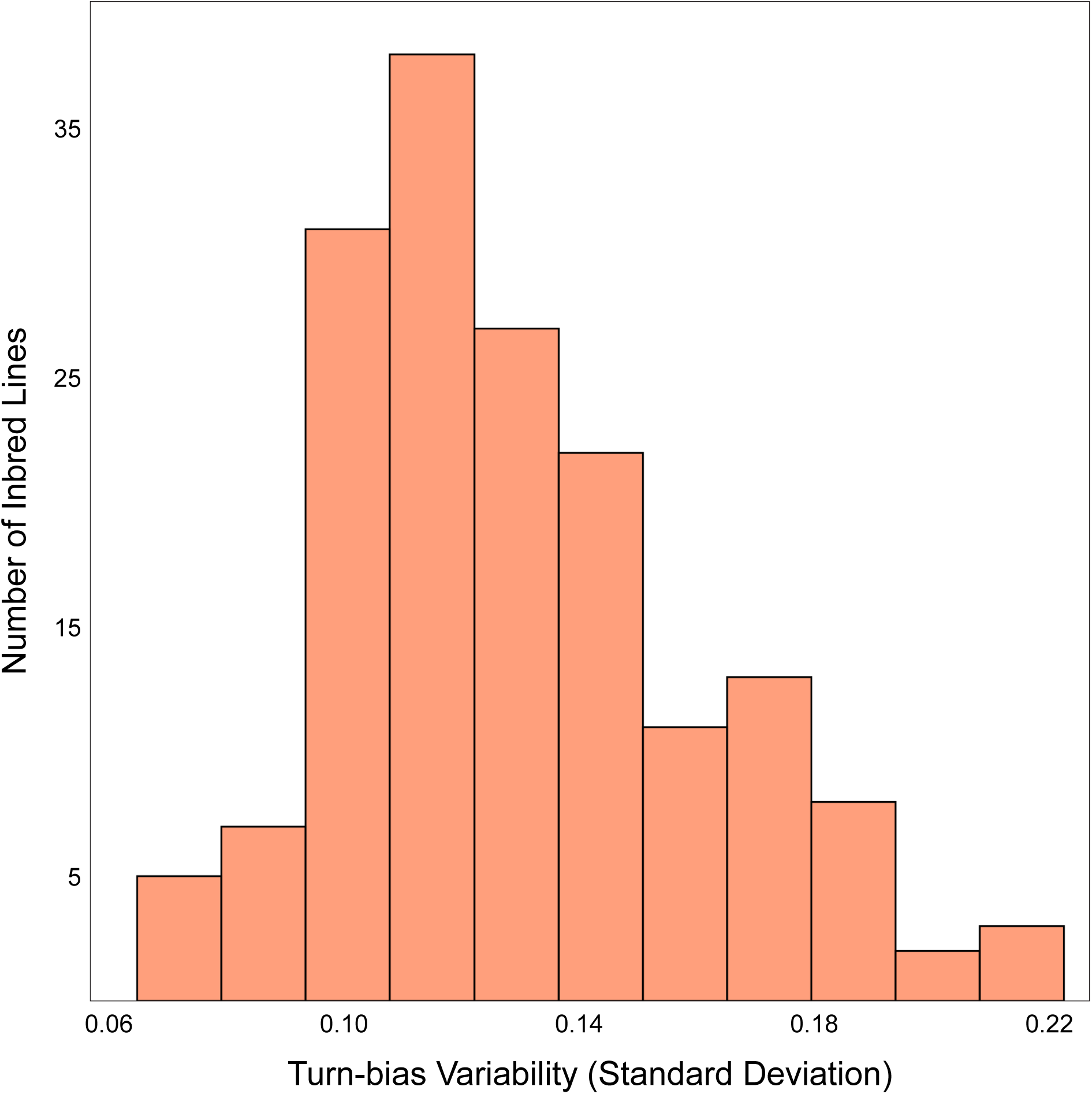
Empirical turn bias variability scores. Histogram of turn bias variability as measured across 167 inbred fruit fly lines [1]. We mapped the summed variant effect sizes (and epistatic effects, if they were included in a model run) of individuals in our simulation to this distribution to determine the latent turn bias variability from which their individual turn bias phenotypes were drawn.

**Figure S3:**
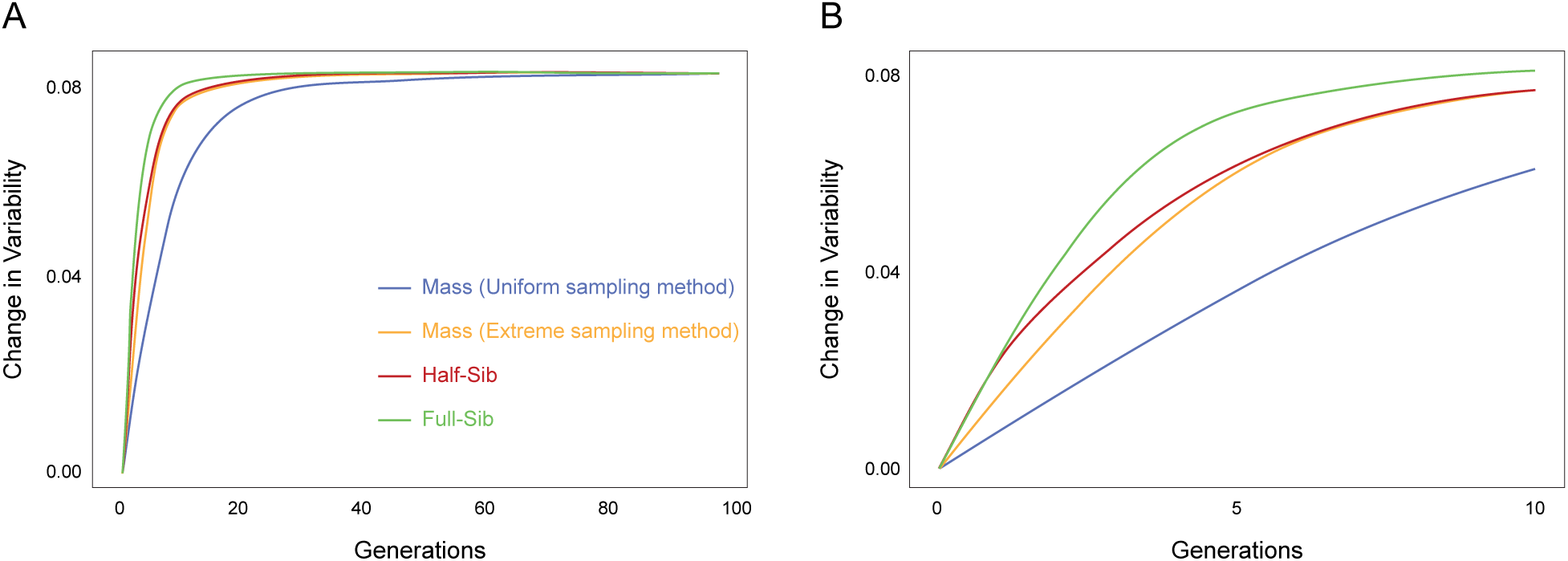
Comparison of mass selection methods. (A) Average change in variability as a function of the number of generations of selection for different selection regimes, including mass selection implemented by keeping individuals with most extreme phenotypic values; (B) Average change in variability for the first 10 generations of selection

**Figure S4:**
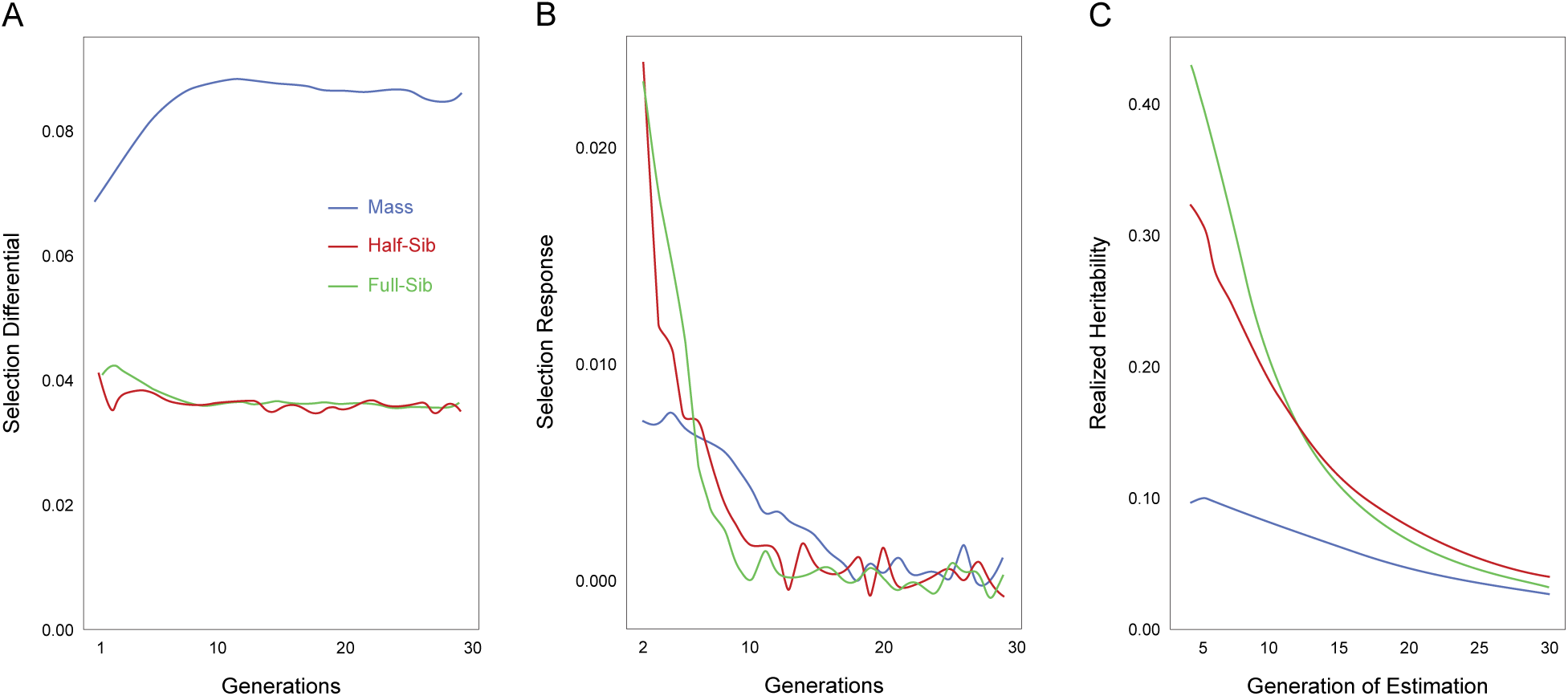
Selection differential, selection response and realized heritability across selection regimes. (A) Average value of selection differential *S* for the first 30 generations of selection, averaged over 100 model runs; (B) Average selection response *R* for the first 30 generations of selection; (C) Estimates of realized heritability *h*^2^ at different time points across regimes

**Figure S5:**
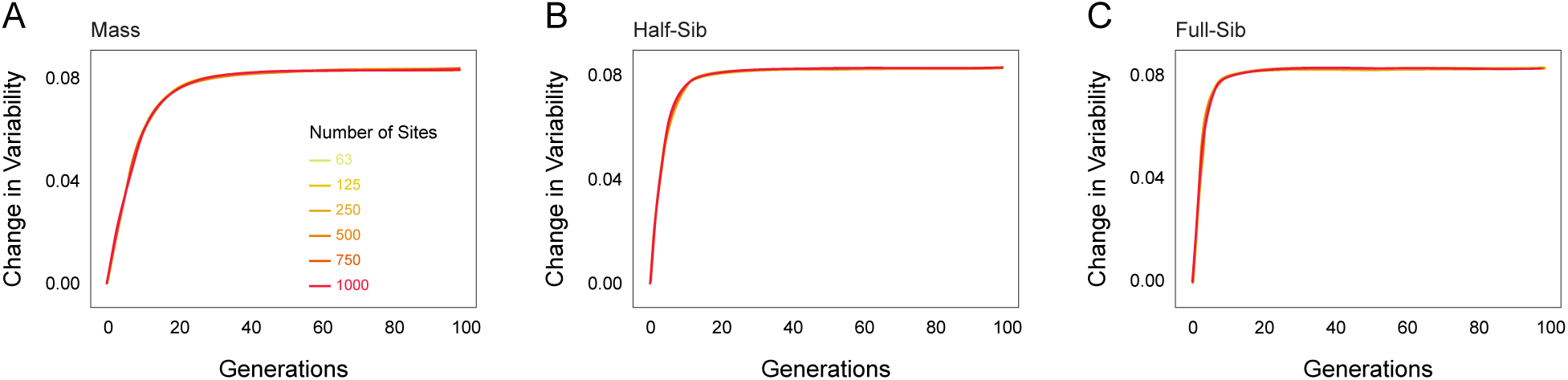
Effect of number of sites on selection response. Average change in variability as a function of the number of generations of selection with varying number of sites that contribute to the variability phenotype in (A) Mass Selection; (B) Half-Sib Selection and (C) Full-Sib Selection

**Figure S6:**
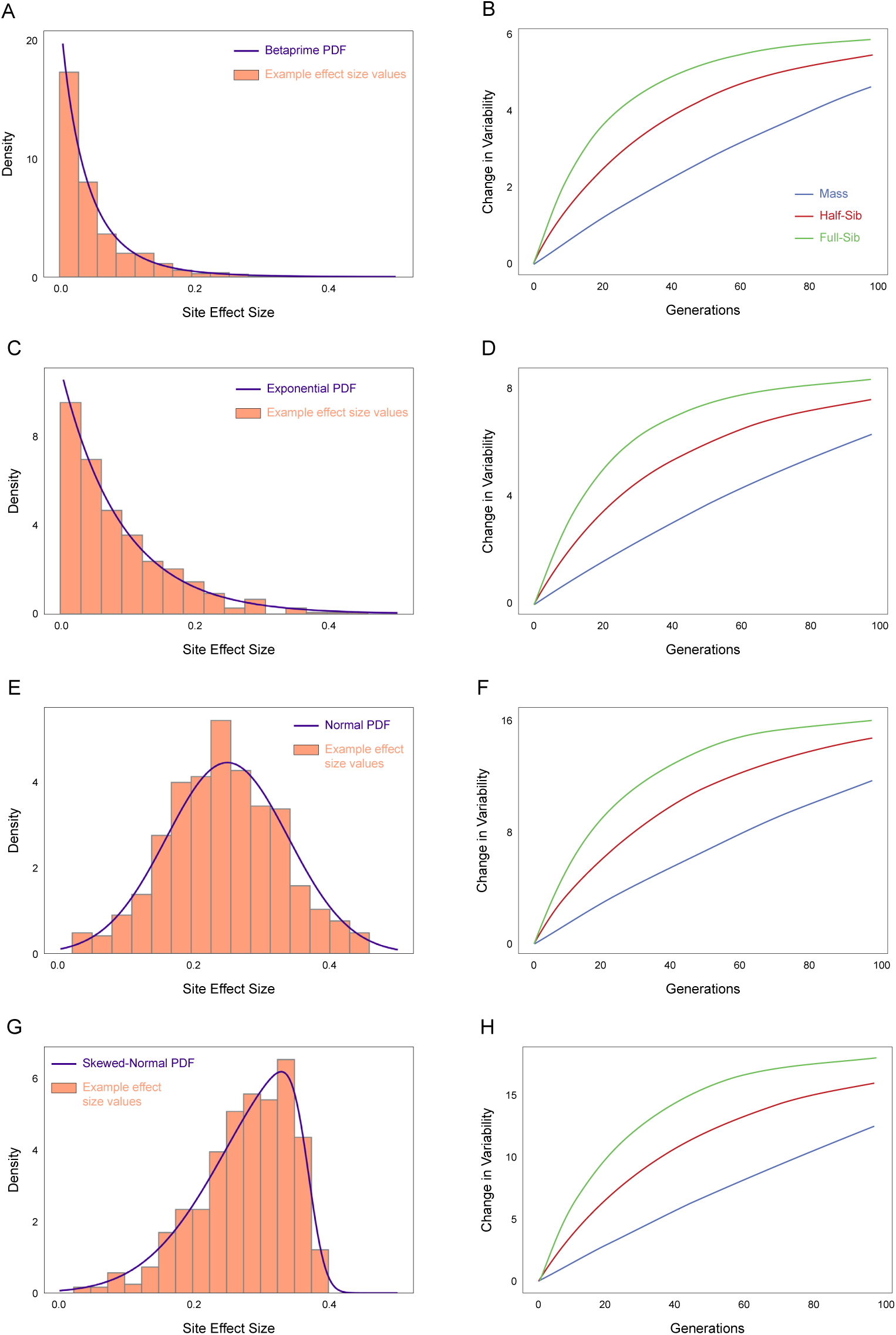
Effect of site effect size distribution on selection response. Probability density functions (PDFs) of distributions and corresponding example values of effect sizes for sites that contribute to variability for (A) Betaprime distribution, (C) Exponential distribution, (E) Normal distribution and (G) Skewed-Normal distribution; (B,D,F,H) Corresponding change in variability (i.e., standard deviation) as a function of number of generations of selection for various selection regimes. [(*f* (*G_a_*) = *G_a_*) for all cases]

**Figure S7:**
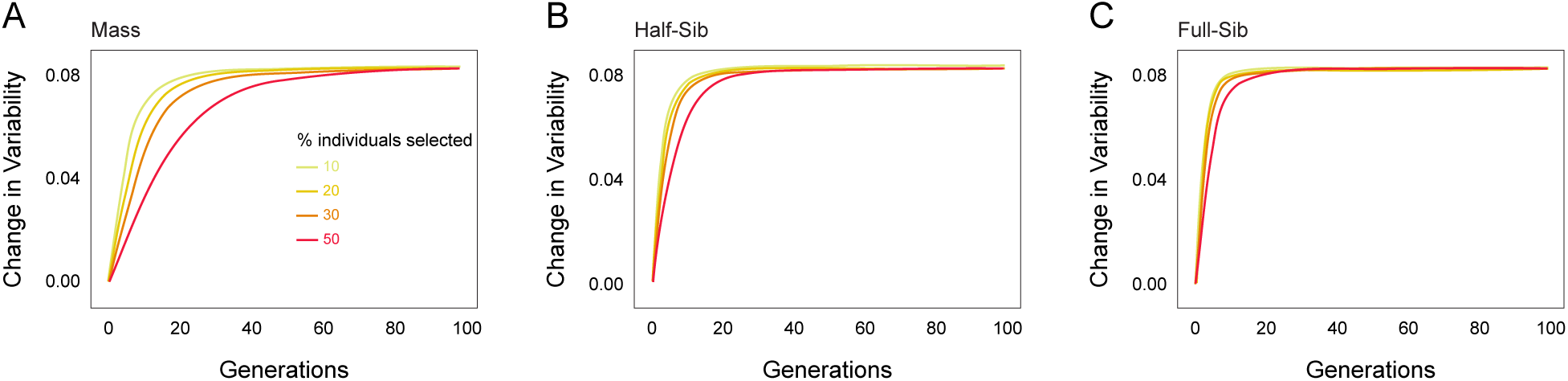
Effect of selection strength on selection response. Average change in variability as a function of the number of generations of selection with varying percentage of individuals kept every generation to sire offspring in (A) Mass Selection; (B) Half-Sib Selection and (C) Full-Sib Selection

**Figure S8:**
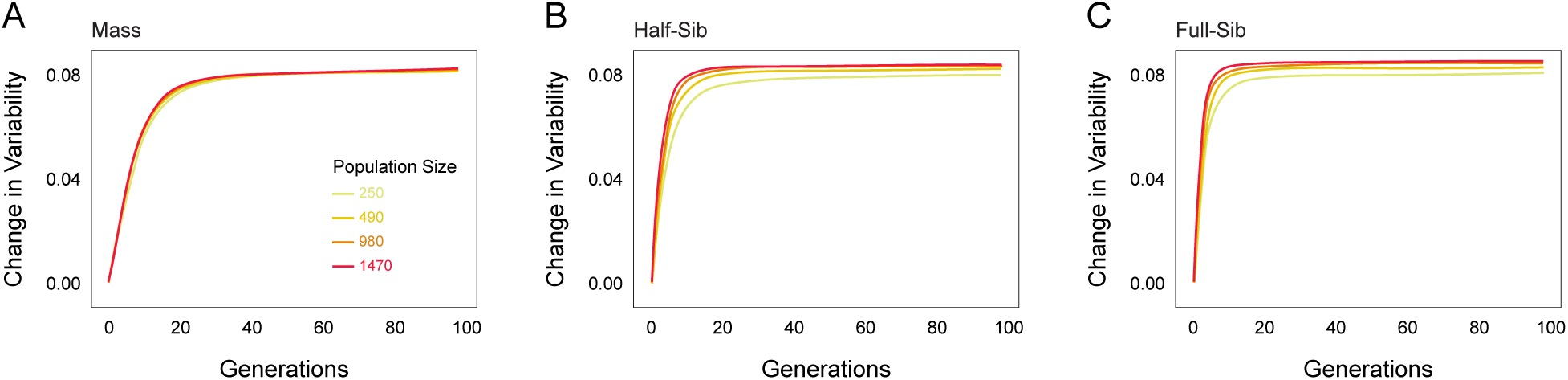
Effect of population size on selection response. Average change in variability as a function of the number of generations of selection with varying total population size in (A) Mass Selection; (B) Half-Sib Selection and (C) Full-Sib Selection

**Figure S9:**
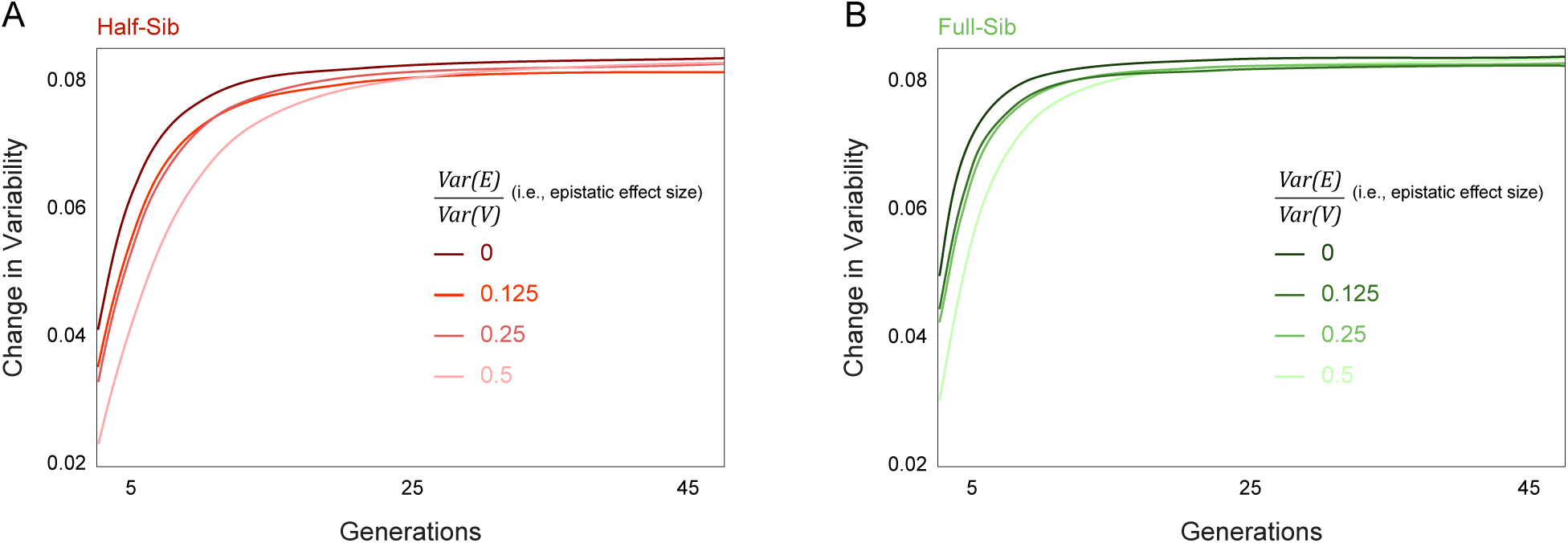
Effect of epistasis on family-based selection response. Average change in variability as a function of the number of generations of selection with varying contributions of epistatic interactions on variability phenotype in (A) Half-Sib Selection and (B) Full-Sib Selection

**Figure S10:**
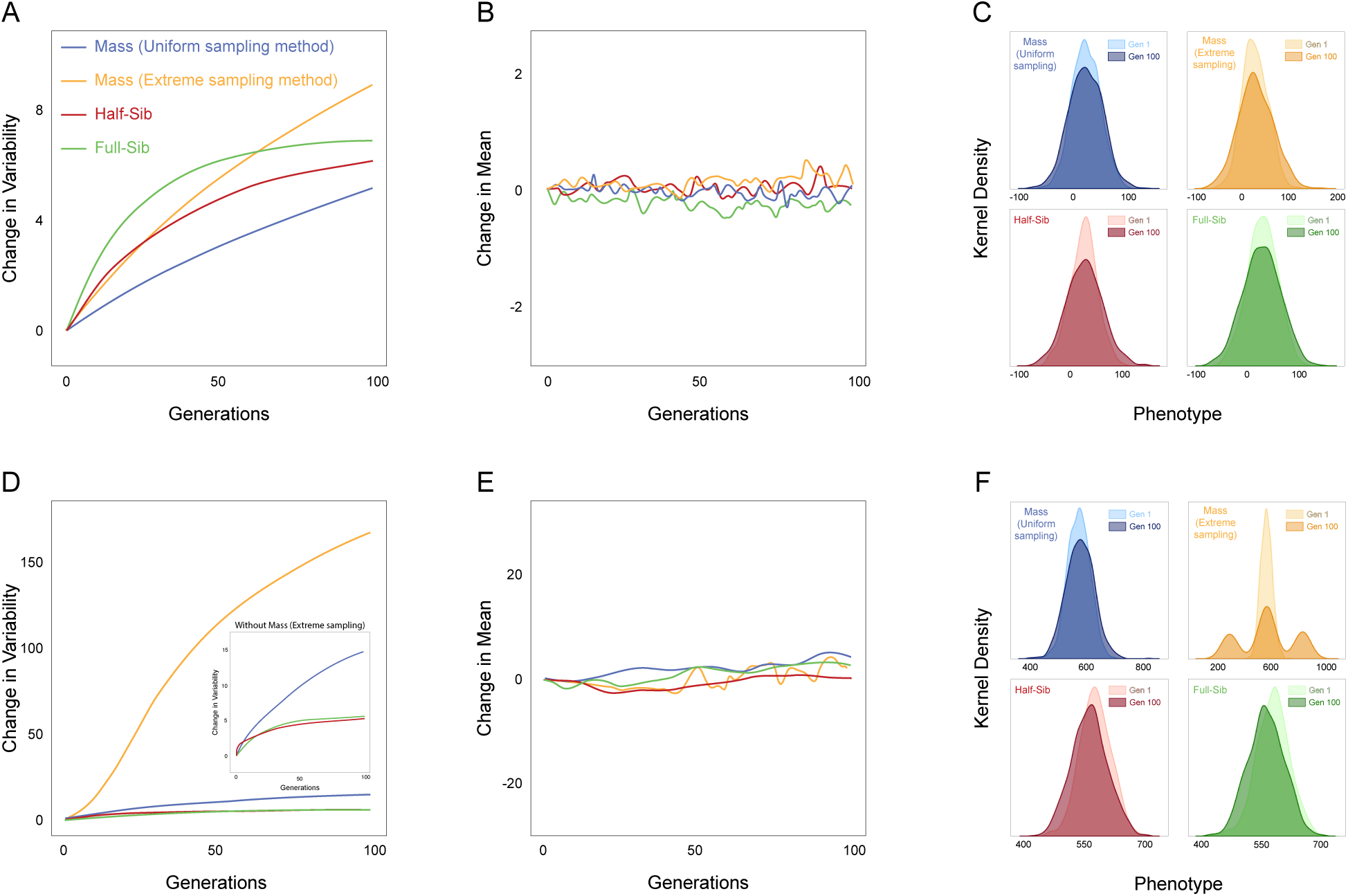
Selection response when both mean and variability are heritable. Panels (A-C) assume trait mean and variability have independent sites with effects drawn from identical distributions. Panels (D-F) assume 15x larger site effects for the mean of the trait versus its variability. A,D) Average change in variability as a function of the number of generations of selection. Inset in D magnifies the responses to selection regimes other than extreme mass selection. (B,E) Average change in mean as a function of number of generations of variability selection. (C,F) Example kernel density estimates of the distributions of trait values at generations 1 and 100 across selection regimes

## Notes

https://lab.debivort.org/family-selection-for-variability/

### Competing Interest Statement

The authors have declared no competing interest.

### Summary of Updates

Introduction and discussion updated to contextualize the paper within the quantitative genetics field; additional supplementary figures, results and discussion added to generalize the model to more traits

